# Systemic metabolic changes in acute and chronic lymphocytic choriomeningitis virus infection

**DOI:** 10.1101/2024.08.09.607318

**Authors:** Caroline Bartman, Shengqi Hou, Fabian Correa, Yihui Shen, Victoria da Silva-Diz, Maya Aleksandrova, Daniel Herranz, Josh Rabinowitz, Andrew Intlekofer

**Author notes:** Equal contribution (Caroline Bartman, Shengqi Hou).

## Abstract

Viral infection of cells leads to metabolic changes, but how viral infection changes whole-body and tissue metabolism *in vivo* has not been comprehensively studied. In particular, it is unknown how metabolism might be differentially affected by an acute infection that the immune system can successfully clear, compared to a chronic and persistent infection. Here we used metabolomics and isotope tracing to identify metabolic changes in mice infected with acute or chronic forms of lymphocytic choriomeningitis virus (LCMV) for three or eight days. Both types of infection alter metabolite levels in blood and tissues, including itaconate and thymidine. However, we observed more dramatic metabolite changes in the blood and tissues of mice with chronic LCMV infection compared to those with acute infection. Isotope tracing revealed that the contribution of both glucose and glutamine to the tricarboxylic acid (TCA) cycle increase in the spleen, liver, and kidneys of mice infected with chronic LCMV, while acute LCMV only increases the contribution of glutamine to the TCA cycle in the spleen. We found that whole-body turnover of both glutamine and thymidine increase during acute and chronic infection, whereas whole-body glucose turnover was surprisingly unchanged. Activated T cells *in vitro* produce thymidine and mice with T cell leukemia display elevated serum thymidine, nominating T lymphocytes as the source of thymidine in LCMV infection. In sum, we provide comprehensive measurements of whole-body and tissue metabolism in acute and chronic viral infection, and identify altered thymidine metabolism as a marker of viral infection.

## Introduction

Viral infection and the resultant host immune response result in major physiologic changes. The infecting virus replicates thousands of times and spreads from cell to cell, requiring RNA production (and DNA if a DNA virus) and protein synthesis of the viral capsid and nonstructural proteins. Immune cells, which account for around 2% of the mass of the human body in the uninflamed state^1^, proliferate wildly-for example T cells specific for the infecting virus can increase 1000-fold- and produce effector proteins^2,3^. These phenomena suggest that the viral infection and the host immune response may be supported by metabolic changes, to produce energy and provide building blocks to enable protein production and cell proliferation. However, how metabolism changes during viral infection *in vivo* is incompletely understood.

One well-studied model of viral infection is the mouse RNA virus lymphocytic choriomeningitis virus (LCMV). This virus infects most tissues of the mouse, including the spleen, liver, and kidney^4^. It induces a strong CD8+ T cell response, and the immune response itself is responsible for the pathology and tissue damage during the infection^4^. Different strains of LCMV, distinguished by only a few nucleotide changes, can result in dramatically different disease outcomes. The host T cell response successfully clears the Armstrong strain of LCMV, while the Clone 13 strain establishes a chronic infection which the immune system cannot clear, resulting in immune dysfunction and T cell exhaustion^5^.

Previous studies have demonstrated that LCMV infection of mice can change serum and tissue metabolite levels, but there have been few measurements of how metabolic flux may change during viral infection. Both kynurenine^6,7^ and itaconate^8^, two well-studied immunometabolites^9–14^, increase in the serum of LCMV infected mice. Tissue metabolite levels have been less investigated, though one study showed that itaconate and pyrimidine metabolites are elevated in the infected liver^8^. One study suggested that liver urea cycle flux may be increased in LCMV infection^15^. Another study from the same group showed that mouse activity and whole-body oxygen consumption decrease at the peak of infection (days 7-8) in chronic LCMV infection^16^.

These past studies are informative yet leave several key questions unanswered. How do metabolite levels change during infection in tissues other than liver? How does the use of specific nutrients change during viral infection, both at the whole-body level and in specific tissues? Physiologic changes like fasting-feeding^17,18^, feeding a ketogenic diet^19^, and cold exposure^20^ cause dramatic changes to the use of nutrients like glucose, lactate, and glutamine, but it is unknown what may happen during viral infection and immune response. Finally, how does the metabolic response differ between acute and chronic viral infection? Since these two types of infection vary dramatically in viral kinetics, as well as the character of T cells responding, we might expect resulting metabolic differences.

Here, we used infusion of stable-isotope-labeled metabolites to measure whole body and tissue-specific nutrient use during both acute and chronic LCMV infection, as well as metabolomics to measure serum and tissue metabolites. Both acute and chronic infection alter blood and tissue metabolite levels, but chronic infection changes more metabolites to a greater extent. We observed increased whole-body turnover of both glutamine and thymidine, a pyrimidine metabolite used for nucleotide salvage production in acute and chronic infection. Chronic infection caused more dramatic changes in nutrient use in highly-infected tissues: glucose and glutamine contribution to the TCA cycle both increase in spleen, liver, and kidney in chronic LCMV infection, while only spleen glutamine use increases in acute infection. Finally, we propose that thymidine production in LCMV infection may be mediated by T cells, since thymocytes, activated T cells in culture, and T-acute lymphoblastic leukemia all display high thymidine levels. Overall, this work provides an atlas of whole-body and tissue metabolite and metabolic flux changes in acute and chronic LCMV infection, and nominates thymidine levels and turnover as a biomarker of viral infection.

## Results

### Chronic LCMV infection changes serum metabolite levels more dramatically than acute infection

We set out to measure how acute and chronic lymphocytic choriomeningitis virus (LCMV) infections in mice change whole-body and tissue metabolism. Conceptually, measuring metabolite changes in LCMV with metabolomics can nominate pathways with altered metabolic flux in infection, and these fluxes can then be measured using stable-isotope metabolite infusion. We first examined changes in serum metabolite levels in acute and chronic LCMV infection. Since the blood delivers metabolites to all tissues and removes metabolites that tissues release, it can give an overview of whole-body metabolic changes.

We measured serum metabolite levels in mice on the third day of LCMV infection (the peak of viremia in acute infection^3^), mice on the eighth day of infection (the peak of T cell numbers in chronic and acute infection), and in uninfected mice. Many metabolites were increased or decreased in the serum in both types of infection (Figure 1A-B). Many of these metabolites were significantly changed in both datasets, including the immunometabolite itaconate and the pyrimidine nucleoside thymidine. When we examined the levels of itaconate and thymidine in the blood across multiple experiments, we found that both metabolites changed significantly more on day 8 of chronic infection compared to day 8 of acute infection (Figure 1C-D). Itaconate is an immunometabolite produced by activated macrophages^9,10^ which affects macrophage function by a number of proposed mechanisms inducing type I interferon^21^ and reducing IL-6 production^22^. We also observed that several deoxypyrimidines involved in DNA synthesis including thymidine, deoxycytidine, and deoxyuridine were increased in both types of infection, while most other nucleoside levels were unchanged (Figure S1A) consistent with a previous report^8^.

**Figure 1:**
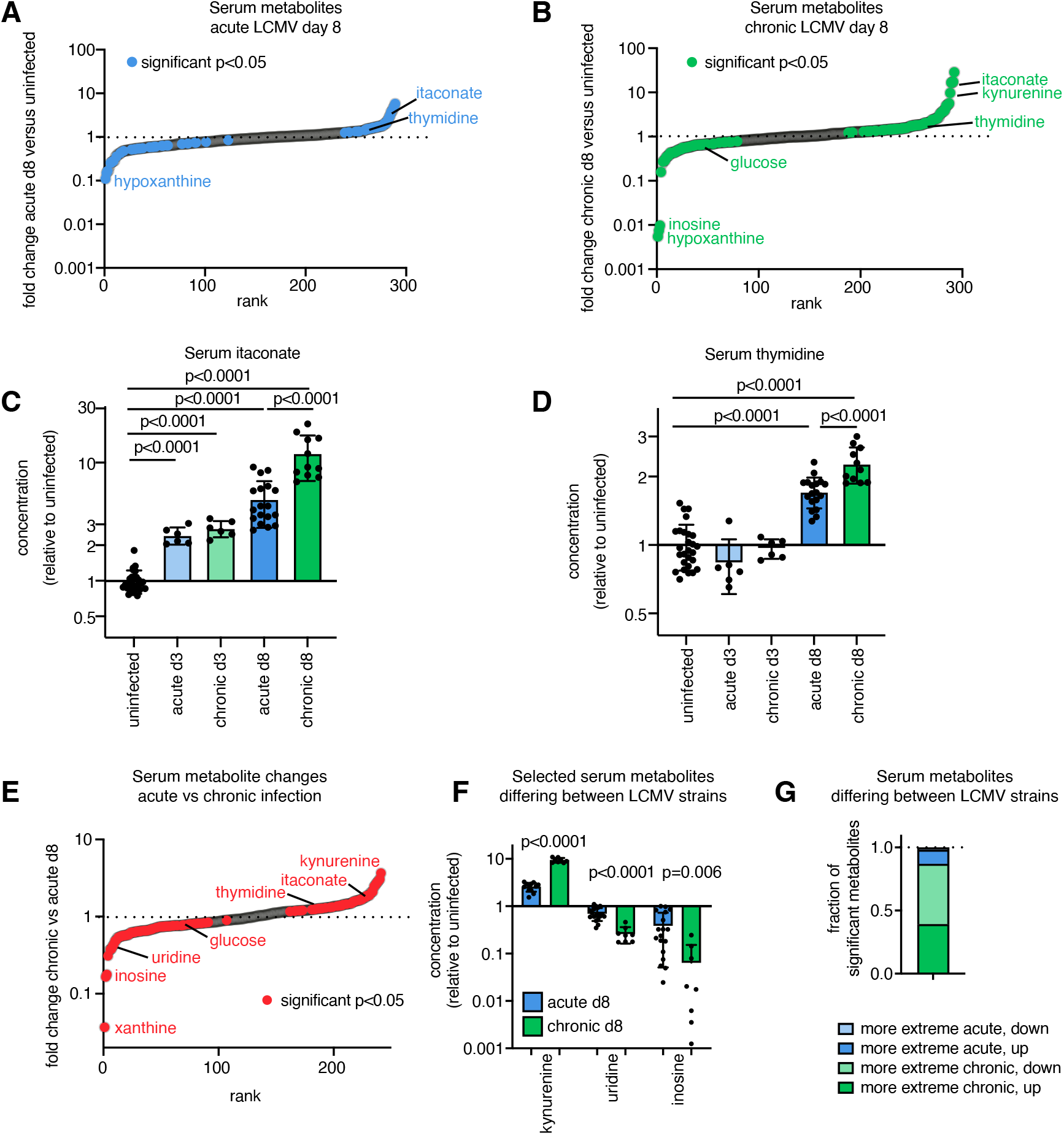
Chronic LCMV infection changes serum metabolite levels more dramatically than acute infection. (a-b) Serum metabolite changes in day 8 acute Armstrong strain LCMV infection (a) or chronic Clone 13 strain (b) compared to uninfected; n=4 uninfected, n=6 infected for each. Two-tailed t-tests in (a-b) performed on log2 transformed metabolite intensities. (c-d) Serum metabolite levels of itaconate (c) and thymidine (d) in LCMV infected mice. Includes multiple experiments, all infected sera normalized to uninfected sera from the same experiment; n=27 uninfected, n=6 acute day 3, n=6 chronic day 3, n=18 acute day 8, n=11 chronic day 8; two-tailed t-tests in performed on log2 fold changes from uninfected, log2 fold changes between chronic and acute d3, and log2 fold changes between chronic and acute d8. (e-f) All (e) and selected (f) serum metabolite changes in chronic versus acute day 8 infection, ratios of fold changes compared to uninfected shown. (g) Summary of significant serum metabolite differences from uninfected on day 8 of acute or chronic infection. For (e-g,) n=18 acute day 8, n=8 chronic day 8, t-tests done comparing log2 fold changes from uninfected.

We next directly compared blood metabolite changes observed on day 8 of acute and chronic LCMV infection, to test whether blood metabolite levels could distinguish these two types of infection. Since we did not conduct infections with both strains in the same experiment, for each experiment we normalized each metabolite to its level in uninfected mice in the same experiment, then compared metabolite fold changes across three independent experiments of acute infection and two of chronic infection. We found that a number of metabolites increase more in chronic infection, including kynurenine^6^, itaconate, and thymidine (Figure 1C-F), while both uridine and inosine decrease more in chronic than acute infection (Figure 1E-F, Figure S1A-B). In general, chronic LCMV infection causes more extreme metabolite increases and decreases across the board (Figure 1G). Of all metabolites significantly different in the blood between acute and chronic infection, 87% were changed to a greater extent in chronic infection.

### Acute LCMV alters tissue metabolites

We next measured which metabolites are altered in tissues on day 8 of acute LCMV infection. The spleen, liver, and kidney are all sites of LCMV infection^4^, while the spleen also serves as a main site for mounting the adaptive immune response to LCMV, so we expected that metabolite levels might be changed by infection in these tissues. Using principal component analysis, infected spleens, livers, and kidneys could be separated from their uninfected counterparts (Figure 2A), although unsurprisingly, tissue identity caused more dramatic metabolite differences than infection status (Figure 2A-B). Certain metabolites in each tissue change dramatically with acute LCMV infection (Figure 2C-F). These changes include increased itaconate in spleen, liver, kidney, and intestine and increased thymidine in the liver.

**Figure 2:**
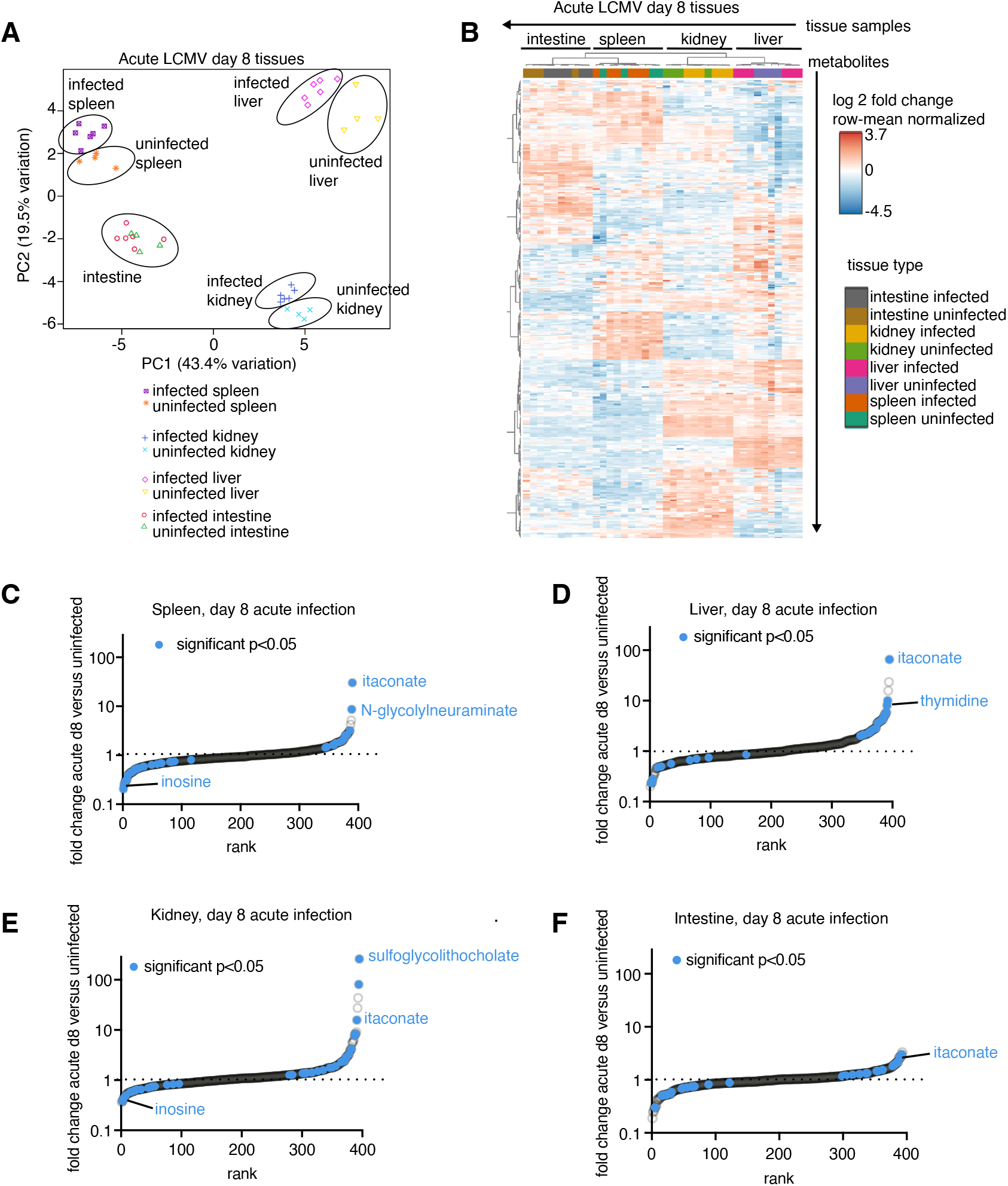
Acute LCMV infection alters tissue metabolites. (a) Principal component analysis of metabolite levels in different tissues of mice on day 8 of acute Armstrong strain LCMV infection or uninfected. Metabolites are log transformed, row-mean normalized, and standard deviation set to 1. Heatmap with hierarchical clustering of metabolite levels, same data as 2a. Metabolites are log transformed and row-mean normalized. (c-f) Tissue metabolites in (c) spleen, (d) liver, (e) kidney, (f) small intestine of mice on day 8 of acute LCMV infection compared to uninfected. Selected metabolites significantly changed from uninfected tissue are labeled. In all panels, n=4 mice uninfected, n=6 infected. T tests in c-f performed on log2 transformed metabolite intensities.

### Chronic LCMV infection changes tissue metabolites more than acute infection

We next asked whether tissue metabolite levels change on day 8 of chronic LCMV infection. We observed a number of metabolite changes in the spleen and liver, including increased itaconate and kynurenine in the spleen (Figure 3A), and increased itaconate and thymidine in the liver (Figure 3B). However, unlike in the blood, tissue thymidine and itaconate changed similarly in acute and chronic LCMV infection (Figure 3C-F); there were no significant differences between spleen or liver levels of these metabolites between the two forms of infection.

**Figure 3:**
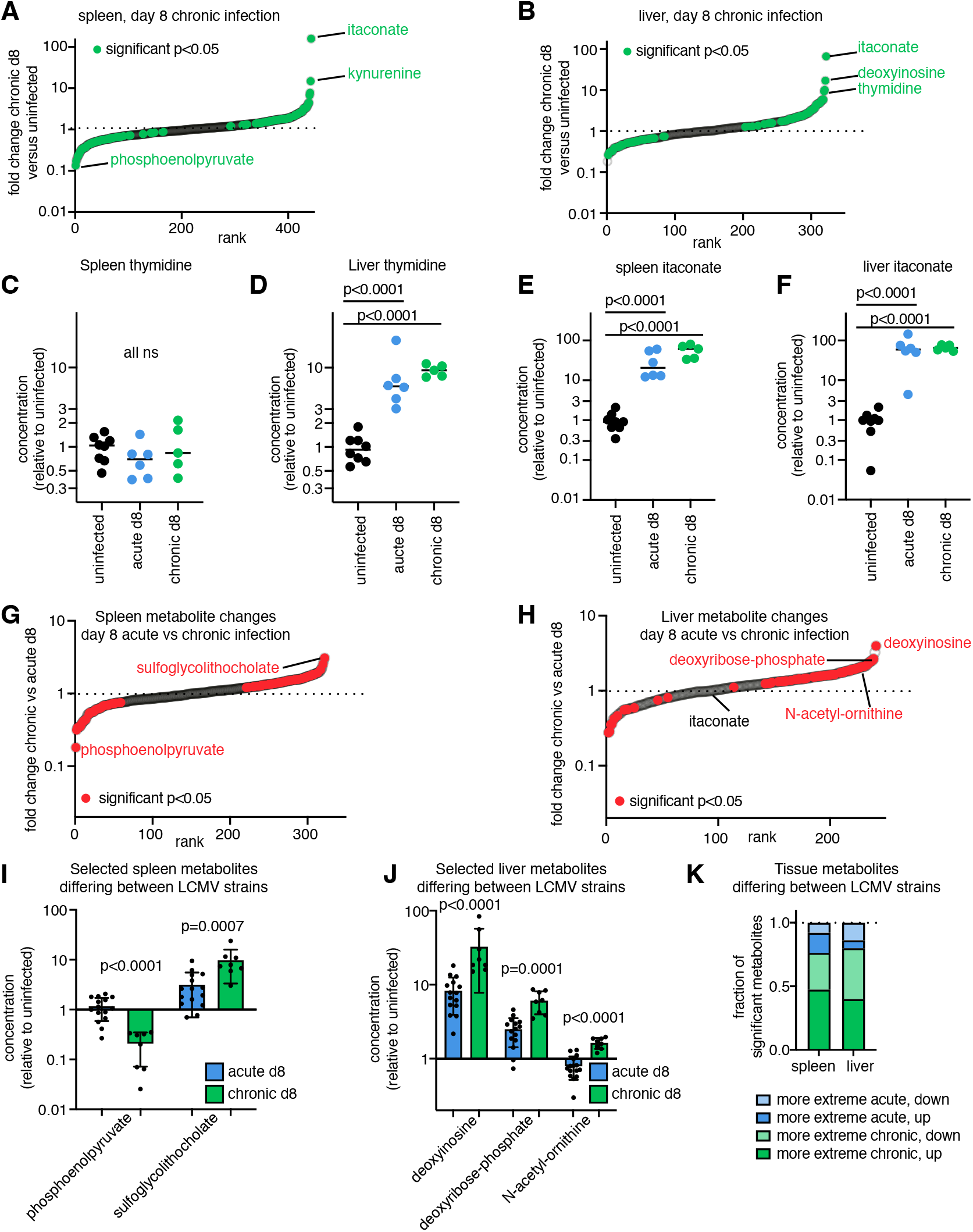
Chronic LCMV infection changes tissue metabolites more than acute infection. Tissue metabolites in (a) spleen or (b) liver of mice on day 8 of chronic infection compared to uninfected; n=4 uninfected, n=5 infected. Two-tailed T-tests in (a-b) performed on log2 transformed metabolite intensities. (c-f) Levels of (c) spleen or (d) liver thymidine, and (e) spleen and (f) liver itaconate in uninfected mice or mice on day 8 of LCMV infection. N=8 uninfected, n=6 acute LCMV, n=5 chronic LCMV; infected values normalized to uninfected mice from the same experiment. T-tests in (c-f) performed on log2 fold changes compared to uninfected from each experiment. (g-h) Spleen (g) and liver (h) metabolite changes from uninfected on day 8 of chronic or acute infection. (i-j) Selected spleen (i) and liver (j) metabolite changes on day 8 of acute or chronic infection, relative to uninfected serum. (k) Summary of significant tissue metabolite differences from uninfected on day 8 of acute or chronic infection. For (g-k,) n=15 acute day 8, n=8 chronic day 8, two-tailed t-tests performed on log2 fold changes from uninfected.

We then compared fold-changes of metabolites in tissues on day 8 of chronic versus acute LCMV infection. Though itaconate and thymidine were not changed between the infection states, certain other tissue metabolites were significantly different between the forms of infection. Examples include splenic sulfoglycolithocholate (a bile salt) and liver deoxyinosine, both of which increase more in chronic than acute infection (Figure 3G-J). As in the blood, tissue metabolites tended to display more extreme changes in chronic than acute infection (Figure 3K), with around 80% of metabolites that differed between the LCMV strains showing more dramatic fold-changes in chronic infection.

### LCMV infection increases glutamine and thymidine whole-body turnover

We next measured whole-body turnover of specific metabolites in acute and chronic LCMV infection. Glucose, glutamine, and lactate are three of the nutrients with the highest whole-body turnover and are some of the dominant substrates for ATP generation in tissues ^17,19,23,24^. We hypothesized that viral infection and immune response might change tissue energy and nutrient requirements, and thus might change turnover of these nutrients. Indeed, we had observed that blood glucose level decreases on day 8 of chronic viral infection (Figure 1B, 0.65x of levels in uninfected mice), and a previous report suggested that carbohydrate burning is reduced in chronic LCMV infection^16^. Therefore we asked whether turnover of these nutrients might be altered in chronic or acute viral infection.

To measure nutrient turnover, we infused carbon-13 labeled nutrient intravenously in infected or uninfected mice for 2.5 hours, and sampled blood from the submandibular vein (Figure 4A). Measuring the blood enrichment of the fully labeled carbon-13 nutrient infused allows calculation of that nutrient’s turnover (also known as rate of appearance or F_circ_) using the equation

**Figure 4:**
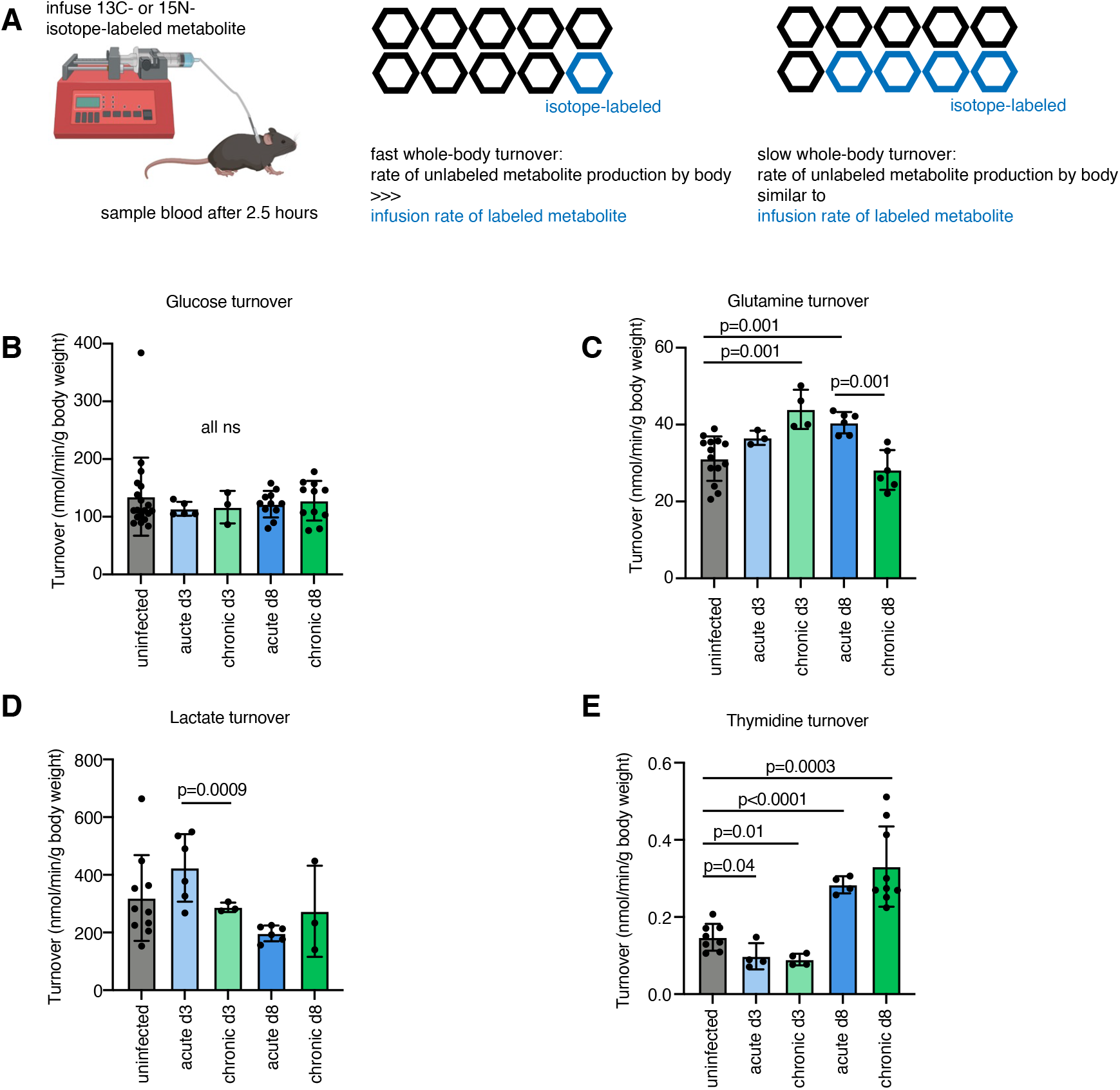
LCMV infection increases glutamine and thymidine whole-body turnover. (a) Schematic of whole-body metabolite turnover measurement using isotope-labeled metabolite infusion. (b) Whole-body turnover of glucose determined by [U-13C] glucose infusion. (5 experiments pooled, overall n=19 uninfected, n=5 acute LCMV day 3, n=3 chronic LCMV day 3, n=11 acute LCMV day 8, n=11 chronic LCMV day 8.) (c) Whole-body turnover of glutamine determined by [U-13C] glutamine infusion. (3 experiments pooled, overall n=14 uninfected, n=3 acute LCMV day 3, n=4 chronic LCMV day 3, n=6 acute LCMV day 8, n=6 chronic LCMV day 8.) (d) Whole-body turnover of lactate determined by [U-13C] lactate infusion. (4 experiments pooled, overall n=10 uninfected, n=6 acute LCMV day 3, n=3 chronic LCMV day 3, n=6 acute LCMV day 8, n=3 chronic LCMV day 8.) (e) Whole-body turnover of thymidine determined by [15N2] thymidine infusion. (2 experiments pooled, overall n=8 uninfected, n=4 acute LCMV day 3, n=4 chronic LCMV day 3, n=4 acute LCMV day 8, n=9 chronic LCMV day 8). All t-tests are Student’s two-tailed t-tests between uninfected vs other groups, acute d3 vs chronic d3, or acute d8 vs chronic d8; all tests not significant if not shown.

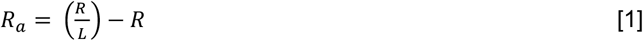

where *R* is rate of carbon-13 nutrient infusion in nanomoles per minute per gram body weight, and *L* is enrichment of fully-labeled nutrient measured in the blood^17,25,26^. Surprisingly, we found that glucose turnover does not change on day 3 or day 8 of acute or chronic infection (Figure 4B). This observation underlines the importance of measuring nutrient turnover directly, rather than using nutrient level as a proxy for metabolic flux. Glutamine turnover increases significantly on day 3 of chronic infection and day 8 of acute infection (Figure 4C, 1.4x and 1.3x of uninfected turnover respectively). Lactate turnover does not change in infected compared to uninfected mice (Figure 4D). Therefore, turnover of major carbohydrate nutrients did not change on day 3 or 8 of LCMV infection compared to uninfected mice, but glutamine turnover was increased.

Next, we measure turnover of thymidine, levels of which increase in the blood and liver in acute and chronic infection (Figures 1 and 3).Thymidine is not a major substrate for energy production; rather, it is a breakdown product of deoxynucleotide synthesis, which can be recycled back into deoxynucleotide production using thymidine kinase or broken down and excreted^27,28^. We found that indeed thymidine turnover doubled on day 8 of both acute and chronic infection (Figure 4E, 1.9x and 2.2x increase relative to uninfected turnover respectively). Therefore, both glutamine and thymidine turnover increased in LCMV infection.

### Glucose contribution to tissue TCA cycle increases in chronic LCMV infection

Carbon-13 nutrient infusion can also reveal which blood nutrients serve as a source for tissue metabolites. For example, the tricarboxylic acid (TCA) cycle, which gives rise to the majority of ATP in all tissues^23^, can be fueled by glucose, glutamine, lactate, fatty acids, and other nutrients. Though whole-body turnover of glucose and glutamine was largely unchanged by LCMV infection, we asked whether their contribution to the TCA cycle of specific tissues might be changed in viral infection.

By infusing carbon-13 glucose and measuring the carbon-13 labeling of tissue malate (Figure 5A), we found that glucose contribution to TCA cycle metabolite increases on day 8 of chronic but not acute LCMV infection in spleen, liver and kidney (Figure 5B-D). The contribution of glucose to liver TCA cycle was also increased on day 3 of chronic but not acute infection (Figure 5C). Some portion of this contribution from glucose is via circulating lactate converted from glucose in other tissues^17^, but overall infection did not increase the contribution from [U-^13^C] lactate itself (Figure S2-no change in spleen or kidney, lactate contribution decreased in liver on day 8 of acute infection). Contribution of glucose to serine and glycine in the spleen was increased, similar to the increase in TCA cycle contribution of glucose, on day 8 of chronic infection (Figure S3). However, contribution of glucose to quadriceps muscle was unchanged on day 8 of chronic LCMV infection (Figure 5E). Muscle represents the most massive tissue type in the body, accounting for about 40% of mouse body weight^29^. Therefore it appears that chronic-infected tissues increase their glucose use, but since muscle (and perhaps other tissues) do not increase glucose use, whole body glucose turnover is unchanged (Figure 4).

**Figure 5:**
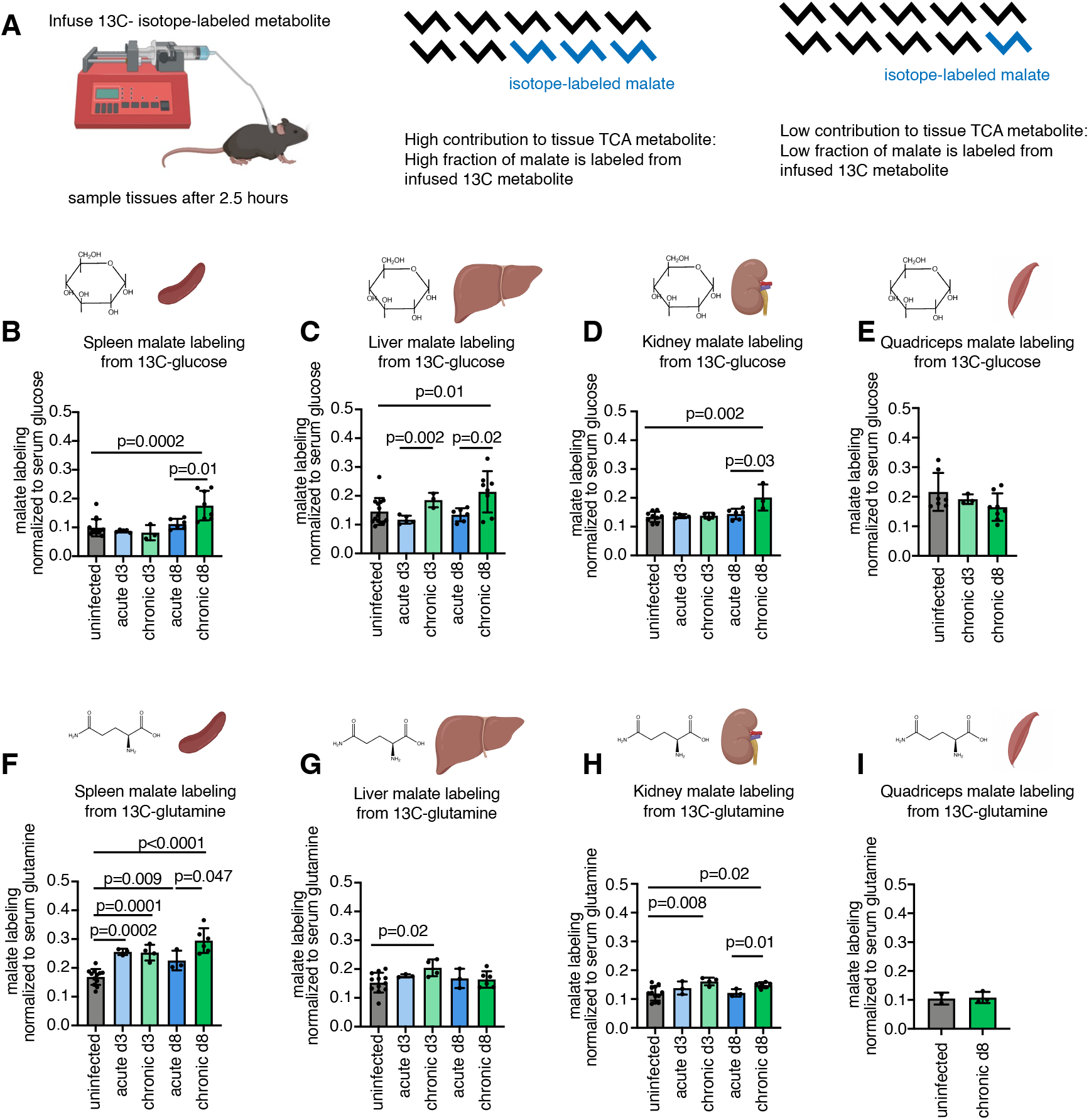
Glucose contribution to tissue TCA cycle increases in chronic LCMV infection. (a) Schematic of measurement of nutrient contribution to TCA metabolites using labeled nutrient infusion. (b) Average carbon labeling of malate TCA intermediate in spleen from [U-13C] glucose infusion in LCMV-infected and uninfected mice. (N=14 uninfected, n=5 acute LCMV day 3, n=3 chronic LCMV day 3, n=6 acute LCMV day 8, n=8 chronic LCMV day 8.) (c) Average labeling of malate in liver from [U-13C] glucose infusion. (N=14 uninfected, n=5 acute LCMV day 3, n=3 chronic LCMV day 3, n=6 acute LCMV day 8, n=8 chronic LCMV day 8.) (d) Average labeling of malate in kidney from [U-13C] glucose infusion. (N=9 uninfected, n=5 acute LCMV day 3, n=3 chronic LCMV day 3, n=6 acute LCMV day 8, n=3 chronic LCMV day 8.) (e) Average labeling of malate in quadriceps from [U-13C] glucose infusion. (N=7 uninfected, n=3 chronic LCMV day 3, n=8 acute LCMV day 8.) (f) Average labeling of malate in spleen from [U-13C] glutamine infusion. (g) Average labeling of malate in liver from [U-13C] glutamine infusion. (h) Average labeling of malate in kidney from [U-13C] glutamine infusion For f-h, n=12 uninfected, n=3 acute LCMV day 3, n=4 chronic LCMV day 3, n=3 acute LCMV day 8, n=6 chronic LCMV day 8. (i) Average labeling of malate in quadriceps muscle from [U-13C] glutamine infusion; n=2 uninfected, n=3 chronic LCMV day 8. All t-tests are Student’s two-tailed t-tests between uninfected vs other groups, acute d3 vs chronic d3, or acute d8 vs chronic d8; all tests not significant if not shown.

Glutamine contribution to the TCA cycle, measured by carbon-13 glutamine infusion, increases in the spleen in all infection conditions (Figure 5F), but with a significantly greater increase on day 8 of chronic infection compared to day 8 of acute infection. Glutamine contribution also increases on day 3 of chronic infection in the liver, and days 3 and 8 of chronic infection in the kidney (Figure 5G-I). In sum, the contribution of glucose and glutamine to the TCA cycle rises in infected tissues (spleen, liver, and kidney) in chronic infection, while in acute infection the only contribution change observed was increased glutamine use in the spleen. Therefore, increased carbon-13 glucose contribution to the TCA cycle in infected tissues distinguishes between infection with these two LCMV strains.

### Thymidine synthesis increases in the spleen in LCMV infection

Since we observed that thymidine whole-body turnover increases in LCMV infection, indicating more whole-body thymidine production and more consumption/excretion (Figure 6A) ^25,26^, we wondered which tissue and cell type might be responsible for the increased production of thymidine. We found that thymidine synthesis rate was increased in the spleen in LCMV infection, particularly in day 8 chronic infection (Figure 6B), by measuring labeling of thymidine from carbon-13 glucose infusion. Note that because nucleotide synthesis is relatively slow, not reaching steady state until around 12 hours of infusion^30^, a 2.5 hour infusion of [U-^13^C]glucose captures thymidine synthesis rate, unlike TCA cycle metabolites which have reached steady-state labeling at this timepoint^26^. Synthesis of thymidine from glucose was not detected in liver, kidney, or intestine, so its production may be specific to the spleen, a major site of the adaptive immune response in LCMV.

**Figure 6:**
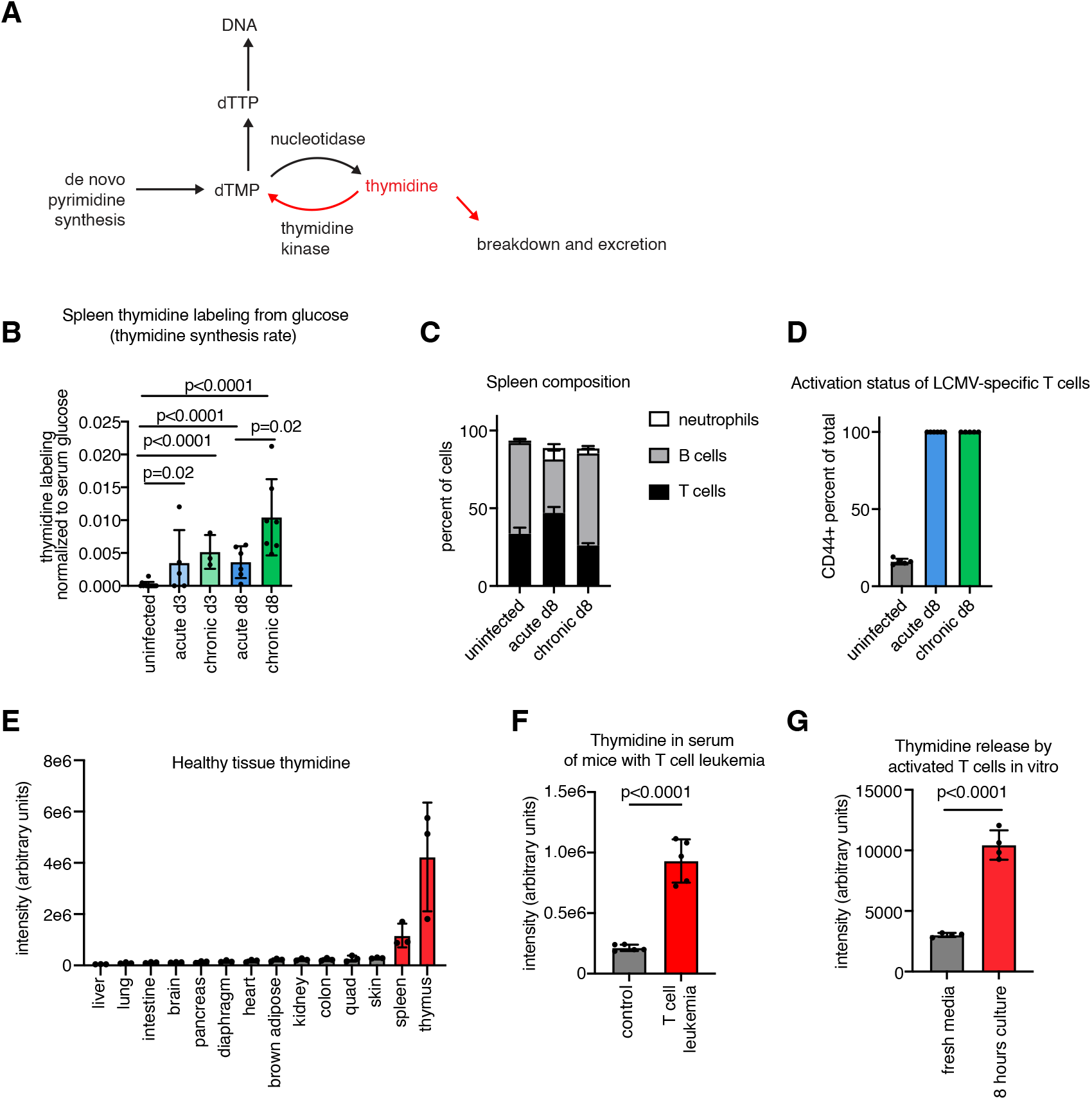
T lymphocytes may be a major source of thymidine. (a) Schematic of thymidine metabolism pathway. (b) Labeling of spleen thymidine from [U-13C] glucose infusion. (5 experiments pooled, overall n=13 uninfected, n=5 acute LCMV day 3, n=3 chronic LCMV day 3, n=6 acute LCMV day 8, n=7 chronic LCMV day 8.) T-tests are Student’s two-tailed t-tests between uninfected vs other groups, acute d3 vs chronic d3, or acute d8 vs chronic d8; all tests not significant if not shown. (c) Fractional composition of CD45+ spleen cells in n=6 uninfected, n=6 day 8 acute, and n=5 day 8 chronic LCMV infected mice measured by flow cytometry. (d) Activation status (CD44+ fraction) of transferred LCMV-specific P14 CD8 T cells in the spleen in n=5 uninfected, n=6 day 8 acute, and n=5 day 8 chronic LCMV infected mice measured by flow cytometry. (e) Thymidine levels in tissues of healthy mice, n=3 mice. (f) Serum thymidine in mice with NOTCH-1 induced T-cell acute lymphocytic leukemia compared to healthy controls, n=5 for each group. (g) Production and release of thymidine by mouse T cells upon activation, n=4 biological replicates per group. All t-tests are Student’s two-tailed t-tests, tests in (f-g) are performed on log2 transformed metabolite intensities.

### T lymphocytes may be a major source of thymidine

We next asked which cells in the spleen might be responsible for producing thymidine during LCMV infection. B and T lymphocytes compose most of the spleen by cell number, both in uninfected and LCMV-infected mice (Figure 6C, not including red blood cells which don’t contain nuclei or most metabolic pathways). However, the proportion of activated and proliferating lymphocytes increases during LCMV infection (Figure 6D), with associated changes in cellular metabolism, such as an increase in the rate of DNA replication^31–34^.

We hypothesize that activated lymphocytes are the likely cellular source of increased thymidine during LCMV infection. In healthy mice, the tissue exhibiting by far the highest level of thymidine is the thymus (Figure 6E), a tissue primarily composed of T cell precursors. Moreover, mice with T-acute lymphocytic leukemia, in which ∼99% of the splenic nucleated cells are T cell blasts^23^, display an even greater increase in serum thymidine than LCMV-infected mice do, suggesting that these dividing T cell blasts are major producers of thymidine (Figure 6F). Finally, activated T cells in culture release thymidine into media (Figure 6G). Collectively, these data suggest that activated T cells in the spleens of LCMV-infected mice are dominant producers of thymidine, contributing to the high thymidine turnover observed in LCMV infection.

## Discussion

In this study, we used metabolomics and labeled nutrient infusions to identify metabolic changes occurring during acute and chronic LCMV infection. We found that while both forms of infection change metabolite levels and metabolite use in tissues, more dramatic changes were observed in chronic infection with the Clone 13 strain. For example, blood levels of itaconate, kynurenine, thymidine, and uridine all change more in chronic Clone 13 than acute Armstrong strain LCMV infection. Similarly, glucose and glutamine contribution to the TCA cycle in the highly-infected tissues spleen, liver and kidney all increase in chronic infection, while only glutamine use in the spleen increases in acute infection. Whole-body nutrient turnovers were somewhat more consistent between acute and chronic: thymidine turnover was increased in both cases, while glutamine turnover increased on day 3 of chronic and day 8 of acute infection. Our data further suggest that activated T cells may be the producers of the increased thymidine in both acute and chronic LCMV infection.

Overall, we observed that chronic LCMV infection changed metabolism more than acute infection. There are two major differences between these disease states: in chronic infection at day 8 virus persists in blood and tissues, while it is low or absent at that time in acute infection; as a consequence, the innate and adaptive immune response persists in chronic infection, while in acute infection at day 8 the immune response is beginning to resolve^3,35^. It is likely that both the innate and adaptive immune response contribute to the metabolic changes observed: for example, itaconate is a hallmark of innate immune activation of macrophages^9,10^ while our data suggests that lymphocytes may be dominant producers of thymidine. In future, it would be valuable to examine metabolite levels and metabolic fluxes at even later stages of infection, where the outcomes of the two types of infection have diverged yet further. It would also be valuable to use cell-type isolation paired with isotope tracing^36^ or imaging mass spectrometry^37,38^ to identify which cell types are responsible for producing different metabolites, or which cell types alter use of different nutrients in the TCA cycle, in acute and chronic infection.

This work nominates increased thymidine in the blood and thymidine whole-body turnover as biomarkers of viral infection. Thymidine turnover was increased in both acute and chronic infection (Figure 4), though blood level (Figure 1) and spleen synthesis rate (Figure 6) were increased more in chronic infection. Based on our observations, we hypothesize that activated T lymphocytes may be primary producers of thymidine. It may be that thymidine turnover increases during any adaptive immune response, or any state when lymphocyte-like cells are proliferating (for example T cell leukemia, Figure 6). In future, we hope to test this association further in other types of infection or vaccination in both mice and humans. However, thymidine metabolism differs between mice and humans, with serum thymidine levels around 10-fold higher in mice^39^, so this feature of infection might not be shared between the species. We also do not yet know the fate of the thymidine produced during LCMV infection: thymidine can be broken down by the liver and excreted or can be recycled into further deoxynucleotide synthesis (Figure 6A). If the latter fate is dominant, then modulating thymidine levels during an infection or vaccination may modulate the immune response or the formation of immune memory, and it would be valuable to test this in future.

We were surprised that LCMV infection does not change whole-body turnover of glucose. How can we reconcile this with the increases in glucose contribution to the TCA cycle in specific tissues (Figure 5), and the previously reported change in carbohydrate burning during chronic LCMV infection^16^? First, spleen, liver, and kidney are relatively small contributors to whole-body TCA flux in the mouse, summing to around 10% of the whole body total^23^, so the increased contribution of glucose measured would not necessarily lead to changes in whole-body glucose consumption. Second, in the study of Baazim and colleagues^16^, the decrease in carbohydrate burning seemed to be driven by a decreased feeding by chronic-infected mice^40^. Since the standard mouse diet is very high in carbohydrates, reduced food consumption would be expected to lower whole-body carbohydrate burning. However in our studies, all mice were fasted for 5 hours before the start of 2.5-hour labeled nutrient infusions, which may have masked the metabolic effect of infection-reduced feeding.

Overall, pairing isotope-labeled nutrient infusions with metabolomics is a powerful approach to measure changes in whole-body and tissue specific metabolism. These approaches have most comprehensively been used to measure metabolism in the context of diabetes/obesity^41,42^, burn wounds^43^, and in recent years, in cancer^44^. However, as a field we know less about the *in vivo* metabolism of other physiologic and pathologic states including infection and immune response, so this study aims to help fill this gap in the field. In future, similar approaches should be used to measure the metabolism of other physiologic states such as cytokine storm, aging, and neurodegenerative disease. Most importantly, such studies can nominate metabolic pathways specifically changed in different disease states, and then such pathways can be modulated to test whether this can improve disease pathology.

## Methods

### Mice

All animal studies were approved either by the Memorial Sloan Kettering Institutional Animal Care and Use Committee (majority of experiments), the Princeton Institutional Animal Care and Use Committee (healthy tissue thymidine concentration and mouse T cell isolation for *in vitro* culture), or the Rutgers Institutional Animal Care and Use Committee (T-acute lymphocytic leukemia experiment). All experiments were performed in C57Bl/6 mice from Charles River Laboratories. Mice for nutrient infusions were purchased with jugular vein catheters implanted by Charles River Laboratories.

### T-cell acute lymphocytic leukemia model

Generation of NOTCH1-induced mouse primary T-cell acute lymphocytic leukemia and secondary transplantation into sub-lethally irradiated recipients was performed as previously described^45^: 1 × 10^6^ leukemia cells were transplanted from primary recipients into sub-lethally irradiated (4.5 Gy) 9-week-old C57BL/6 secondary recipients, which were pre-catheterized by Charles River Laboratories in the right jugular vein. Animals were monitored for signs of distress or motor function daily. Blood was sampled 9-10 days post leukemia transplantation.

### Lymphocytic choriomeningitis virus infection

Armstrong and Clone 13 strains of LCMV were purchased from the European Virus Archive. On the day prior to infection, 5×10^4^ P14 transgenic CD8^+^ T cells (CD45.1^+^) were delivered intravenously to catheterized mice. Mice were infected by intraperitoneal delivery of 2×10^5^ plaque-forming units (PFU) of LCMV Armstrong or intravenous delivery of 4×10^6^ PFU of LCMV Clone 13.

### Nutrient infusions

Nutrient tracers were diluted as follows: [U-^13^C] glucose (CLM-1396, Cambridge Isotope Laboratories) was diluted to 300 mM in sterile saline; [U-^13^C] lactate (20 w/w% solution, CLM-1579, Cambridge Isotope Laboratories) was diluted to 5% (4x dilution, 435mM final concentration) in water; [U-^13^C] glutamine (CLM-1822, Cambridge Isotope Laboratories) was diluted to 100 mM in sterile saline; [^15^N_2_] thymidine (NLM-3901, Cambridge Isotope Laboratories) was diluted to 1mM in sterile saline. Five hours prior to tracer infusion, mice were fasted by switching to fresh cages without food. Mice were weighed and infusions were begun at a rate of 0.1 microliter per minute per gram body weight. Each infusion was continued for 150 minutes. After 150 minutes, mouse blood was collected by submandibular vein sampling. After animal sacrifice by cervical dislocation, tissues were rapidly collected and freeze-clamped using a liquid nitrogen-cooled Wollenberger clamp. Tissues were stored at -80°C until processed.

### T cell *in vitro* stimulation

Mouse spleens were harvested and pooled as single-cell suspensions by passing through 70-μm cell strainers into RPMI 1640 media. After red blood cell lysis (eBioscience, 00-4300-54), naïve CD8^+^ T cells were purified by magnetic bead separation using mouse naïve CD8a^+^ T Cell Isolation Kit (Miltenyi Biotec, 130-096-543) following vendor instructions. Approximately 2-3 million purified naïve CD8+ T cells can be obtained from each mouse. Cells were cultured in complete RPMI media made with RPMI 1640 (11875119, ThermoFisher), supplemented with 10% FBS, 100 U ml^−1^ penicillin, 100 μg ml^−1^ streptomycin and 55 μM 2-mercaptoethanol. Cells were maintained at 1 million cells per ml in 1ml media in 12-well plates. T cells were stimulated for 24 hours with plate-bound anti-CD3 (10 μg ml^−1^, Bio X Cell, BE0001-1) and anti-CD28 (5 μg ml^−1^, Bio X Cell, BE0015-1) in complete RPMI media supplemented with recombinant IL-2 (100 U ml^−1^, Peprotech, 217-12). At 24 hours after stimulation, media was changed on cells and then was sampled for mass spectrometry at 32 hours after stimulation.

### Blood processing and extraction

Sampled blood was kept on ice for up to 60 min after sampling, then centrifuged at 4°C, 15000 RCF for 10 min. The serum fraction was transferred to another tube and stored at -80°C. For mass spectrometry, 2-3 microliters of serum were extracted in 50 volumes methanol on dry ice, then centrifuged at 4°C, 15000 RCF for 25 min, and then transferred to mass spectrometry vials (Thermo Scientific 200046, caps Thermo Scientific 501313) for measurement.

### Tissue processing and extraction

Throughout processing until extraction, tissues were kept buried in dry ice, or in metal Eppendorf racks surrounded by dry ice. Tissues were ground into powder using the Retsch CryoMill. Tissue powder was weighed (5-20mg in dry-ice-precooled Eppendorf tubes), and tissues were extracted by vortexing in 40x volumes precooled acetonitrile-methanol-water (40%/40%/20% v/v/v), then left on water ice over dry ice for 10 minutes. Then the solution was centrifuged at 4°C, 14000 RCF for 25 min, moved to a new tube, then centrifuged again at 4°C, 14000 RCF for 25 min, to remove any particulates. Then extract was moved to mass spectrometry vials for measurement.

### T cell media processing and extraction

Twenty microliters of media was extracted in 80 microliters methanol at -20C. Samples were centrifuged at 4°C, 14000 RCF for 25 min. Then 30 microliters supernatant was added to 90 microliters precooled acetonitrile-methanol-water (40%/40%/20% v/v/v) and moved to mass spectrometry vials for measurement.

### Mass Spectrometry

Water soluble metabolite measurements were obtained by running samples on the Q Exactive Plus hybrid quadrupole-orbitrap mass spectrometer (Thermo Scientific) coupled with hydrophilic interaction chromatography (HILIC). An XBridge BEH Amide column (150 mm × 2.1 mm, 2.5 μM particle size, Waters, Milford, MA) was used. The gradient was solvent A (95%:5% H2O:acetonitrile with 20 mM ammonium acetate, 20 mM ammonium hydroxide, pH 9.4) and solvent B (100% acetonitrile) 0 min, 90% B; 2 min, 90% B; 3 min, 75%; 7 min, 75% B; 8 min, 70% B, 9 min, 70% B; 10 min, 50% B; 12 min, 50% B; 13 min, 25% B; 14 min, 25% B; 16 min, 0% B, 20.5 min, 0% B; 21 min, 90% B; 25 min, 90% B. The flow rate was 150 μL/min with an injection volume of 10 μL and a column temperature of 25°C. The MS scans were in negative ion mode with a resolution of 140,000 at m/z 200. The automatic gain control (AGC) target was 5 × 10^6^ and the scan range was m/z 75−1000. Xcalibur (Thermo Fisher) was used to collect raw data.

### Flow cytometry

Flow cytometry was performed on single-cell suspensions of splenocytes after red blood cell lysis. Fluorescently labelled antibodies were purchased from Biolegend: anti-CD3 PE (17A2), anti-CD4 BV711 (RM4-5), anti-CD8 BV785 (53-6.7), anti-CD11b BV605 (M1/70), anti-CD19 BV510 (6D5), anti-CD44 FITC (IM7), and anti-Ly6G FITC (1A8); or eBioscience: anti-PD1 PE-Cy7 (J43). Data was collected on a Cytek 5 Laser Aurora and analyzed with FlowJo 10 software.

### Mass spectrometry analysis

All LC-MS data, both raw abundances and abundances of labeled forms, were analyzed by El-Maven v0.5.0.

### Metabolomics analysis

Heatmaps and principal component analysis plots were produced using Metaboanalyst (https://www.metaboanalyst.ca/) using the “Statistical Analysis [one factor]” tool. Metabolomics data were log transformed and row-mean centered. When comparing metabolite levels using t-tests, any metabolite intensities less than 1000 were set to 1000 (approximate limit of detection for this mass spectrometer).

### Labeling analysis

For infusion experiments involving carbon-13 or nitrogen-15 labeling, outputs were corrected for natural ^13^C abundance using the accucor package in R (Version 0.2.3).

Whole body turnover of an infused nutrient was calculated using equation

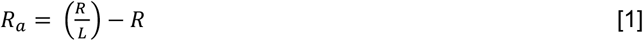

In the case of carbon-13 lactate infusion, all measured L were multiplied by 1.45 before use in equation 1, to correct for the reduced labeling observed when sampling venous blood compared to arterial blood during carbon-13 lactate infusion^17^. (This reduced labeling is likely due to a combination of the high turnover of lactate and the high production of lactate by the tissue bed drained by the vein.)

To calculate the contribution of an infused nutrient to a tissue metabolite, we calculated the average carbon-13 labeling of the infused nutrient in the blood, as well as the average carbon-13 labeling of the nutrient of interest in the tissue, using equation

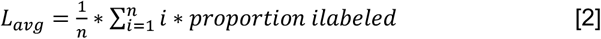

Where *n* is the total number of carbons in the nutrient (i.e. 6 for glucose, 3 for lactate, 5 for glutamine, 4 for malate). Then the contribution of an infused nutrient *x* to a tissue nutrient *y* such as malate is calculated by taking the quotient of *L*_*avg*_ for the tissue nutrient *y* divided by *L*_*avg*_ for the infused blood nutrient *x*,

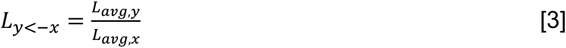

### Statistics and figures

All t-tests are Student’s two-tailed t-tests, and details of experimental sample size and t tests are described in each legend. T-tests on metabolite levels were performed on log-transformed metabolite intensities, except for the comparisons between day 8 of acute and day 8 of chronic infection serum and tissues, in which case t-tests were performed on the log_2_fold change of infected normalized to uninfected. Some schematics were created using BioRender.

---

**Figure S1: Serum nucleoside levels in LCMV infection, related to Figure 1**. (a-b) Serum nucleoside metabolite changes in day 8 acute Armstrong strain LCMV infection (a) or chronic Clone 13 strain (b) compared to uninfected; n=4 uninfected, n=6 infected for each. Two-tailed t-tests in (a-b) performed on log_2_transformed metabolite intensities.

**Figure S2: Lactate contribution to tissue TCA cycle is mostly unchanged by LCMV infection, related to Figure 5**. (a) Average carbon labeling of malate TCA intermediate in spleen from [U-^13^C] lactate infusion in LCMV-infected and uninfected mice. (N=11 uninfected, n=5 acute LCMV day 3, n=3 chronic LCMV day 3, n=6 acute LCMV day 8, n=3 chronic LCMV day 8.) (b) Average labeling of malate in liver from [U-^13^C] lactate infusion. (N=11 uninfected, n=6 acute LCMV day 3, n=3 chronic LCMV day 3, n=6 acute LCMV day 8, n=3 chronic LCMV day 8.) (c) Average labeling of malate in kidney from [U-^13^C] lactate infusion. (N=7 uninfected, n=6 acute LCMV day 3, n=6 acute LCMV day 8.) All t-tests are Student’s two-tailed t-tests between uninfected vs other groups, acute d3 vs chronic d3, or acute d8 vs chronic d8; all tests not significant if not shown.

**Figure S3: Glucose contribution to tissue serine and glycine increases in chronic LCMV infection, related to Figure 5**. (a-b) Average carbon labeling of (a) serine or (b) glycine in spleen from [U-^13^C] glucose infusion in LCMV-infected and uninfected mice. (c-d) Average labeling of (c) serine or (d) glycine in liver from [U-^13^C] glucose infusion. For each graph, n=14 uninfected, n=5 acute LCMV day 3, n=3 chronic LCMV day 3, n=6 acute LCMV day 8, n=8 chronic LCMV day 8.)

