## Supplementary Figures for "Systemic metabolic changes in acute and chronic lymphocytic choriomeningitis virus infection"

### Figure S1

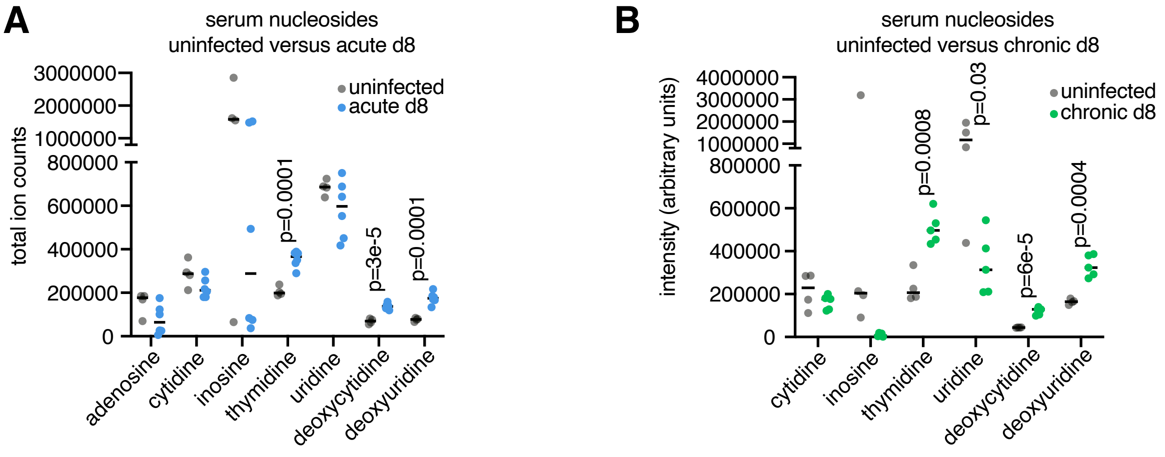


**Figure S1: Serum nucleoside levels in LCMV infection, related to Figure 1.** (a-b) Serum nucleoside metabolite changes on day 8 of acute Armstrong strain LCMV infection (a) or chronic Clone 13 strain (b) compared to uninfected; n=4 uninfected, n=6 infected for each. Two-tailed t-tests in (a-b) performed on log2 transformed metabolite intensities.

### Figure S2

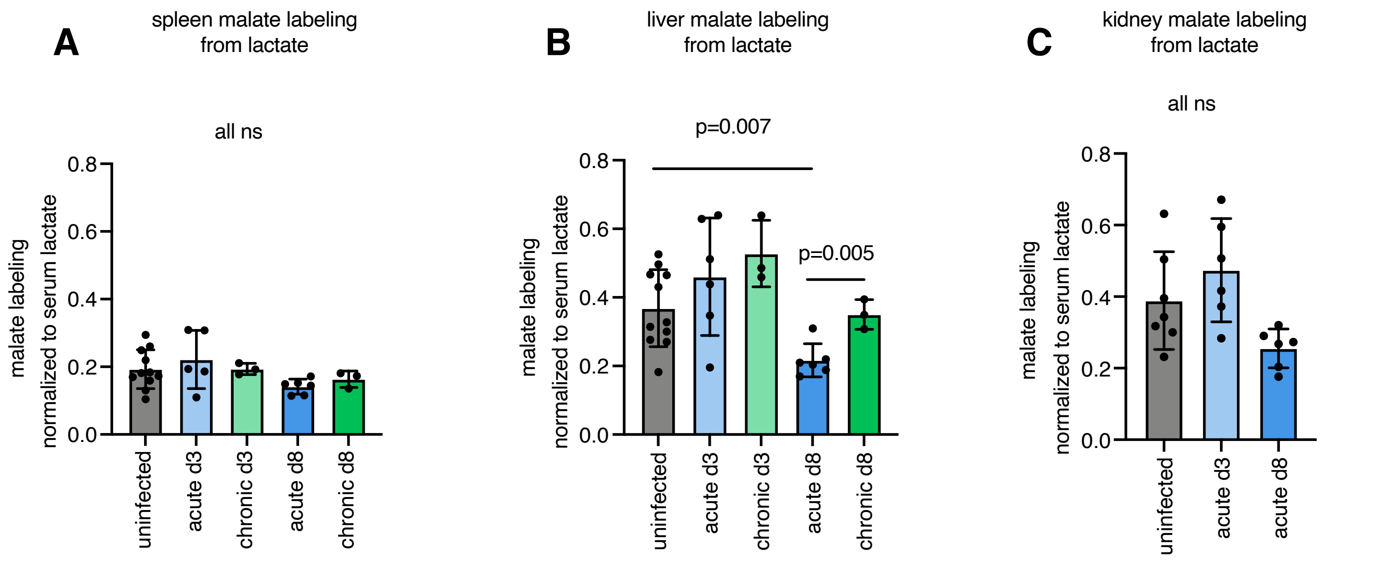


**Figure S2: Lactate contribution to tissue TCA cycle is mostly unchanged by LCMV infection, related to Figure 5.** (a) Average carbon labeling of malate TCA intermediate in spleen from [U-13C] lactate infusion in LCMV-infected and uninfected mice. (N=11 uninfected, n=5 acute LCMV day 3, n=3 chronic LCMV day 3, n=6 acute LCMV day 8, n=3 chronic LCMV day 8.) (b) Average labeling of malate in liver from [U-13C] lactate infusion. (N=11 uninfected, n=6 acute LCMV day 3, n=3 chronic LCMV day 3, n=6 acute LCMV day 8, n=3 chronic LCMV day 8.) (c) Average labeling of malate in kidney from [U-13C] lactate infusion. (N=7 uninfected, n=6 acute LCMV day 3, n=6 acute LCMV day 8.) All t-tests are two-tailed t-tests between uninfected vs other groups, acute d3 vs chronic d3, or acute d8 vs chronic d8; all tests not significant if not shown.

### Figure S3

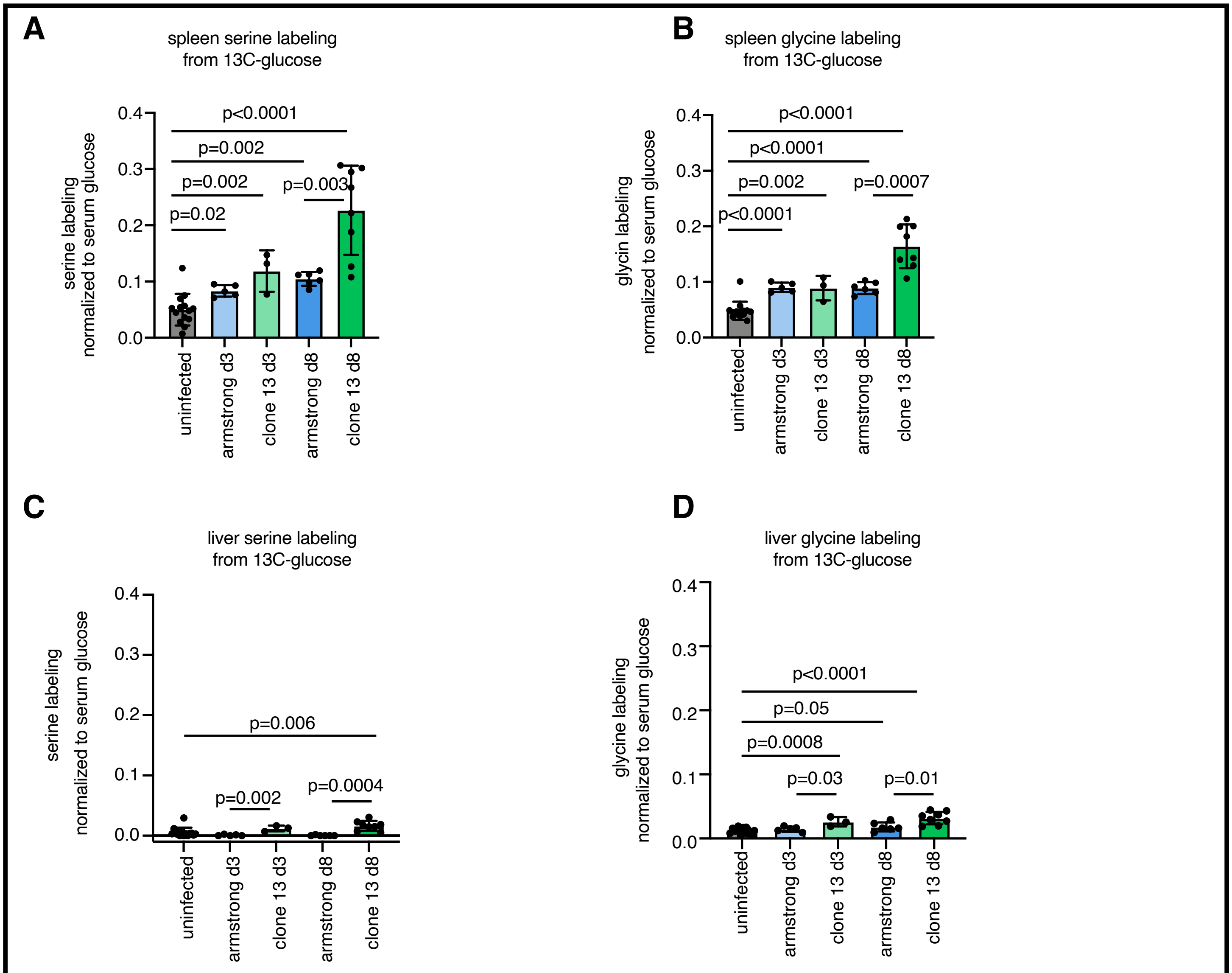


**Figure S3: Glucose contribution to tissue serine and glycine increases in chronic LCMV infection, related to Figure 5.** (a-b) Average carbon labeling of (a) serine or (b) glycine in spleen from [U-13C] glucose infusion in LCMV-infected and uninfected mice. (c-d) Average labeling of (c) serine or (d) glycine in liver from [U-13C] glucose infusion. For each graph, n=14 uninfected, n=5 acute LCMV day 3, n=3 chronic LCMV day 3, n=6 acute LCMV day 8, n=8 chronic LCMV day 8.)
